# Inferring putative transmission clusters with Phydelity

**DOI:** 10.1101/477653

**Authors:** Alvin X. Han, Edyth Parker, Sebastian Maurer-Stroh, Colin A. Russell

## Abstract

Current phylogenetic clustering approaches for identifying pathogen transmission clusters are limited by their dependency on arbitrarily-defined genetic distance thresholds for within-cluster divergence. Incomplete knowledge of a pathogen’s underlying dynamics often reduces the choice of distance threshold to an exploratory, ad-hoc exercise that is difficult to standardise across studies. Phydelity is a new tool for the identification of transmission clusters in pathogen phylogenies. It identifies groups of sequences that are more closely-related than the ensemble distribution of the phylogeny under a statistically-principled and phylogeny-informed framework, without the introduction of arbitrary distance thresholds. Relative to other distance threshold-based and model-based methods, Phydelity outputs clusters with higher purity and lower probability of misclassification in simulated phylogenies. Applying Phydelity to empirical datasets of hepatitis B and C virus infections showed that Phydelity identified clusters with better correspondence to individuals that are more likely to be linked by transmission events relative to other widely-used non-parametric phylogenetic clustering methods without the need for parameter calibration. Phydelity is generalisable to any pathogen and can be used to identify putative direct transmission events. Phydelity is freely available at https://github.com/alvinxhan/Phydelity.

## Introduction

Recent advances in high-throughput sequencing technologies have led to the widespread use of sequence data in infectious disease epidemiology (Gardy and Loman 2017). In particular, epidemiologically relevant information such as the structure of transmission networks and infection source identification are increasingly inferred from virus phylogenies, especially for measurably evolving viral pathogens like HIV-1 and hepatitis C viruses (Ambrosioni et al. 2012; Bezemer et al. 2015; Matsuo et al. 2017; de Oliveira et al. 2017; Charre et al. 2018). Non-parametric phylogenetic-based clustering tools operate on the assumption that pathogens in a transmission cluster are linked by transmission events rapid enough that molecular evolution between the transmitted pathogens is minimal, and thus genetically more similar amongst themselves than to the ensemble of input isolates (Prosperi et al. 2011; Ragonnet-Cronin et al. 2013). This assumption is generally valid for rapidly evolving pathogens such as RNA viruses as genetic changes between sequences sampled from transmission pairs are generally low (Campbell et al. 2018).

Non-parametric phylogenetic clustering methods typically measure the genetic divergence of sequence pairs either by their genetic distances that are computed from the sequence data directly (Aldous et al. 2012; Ragonnet-Cronin et al. 2013) or by their patristic distances from the inferred phylogenetic tree (i.e. the sum of the inferred phylogenetic branch lengths linking the two sequences; Brenner et al. 2007; Prosperi et al. 2011). The divergence of a cluster can be defined as the median (Prosperi et al. 2011) or largest (Ragonnet-Cronin et al. 2013) pairwise distance between member sequences of the cluster. To define transmission clusters, an upper divergence threshold is implemented either as an absolute distance limit (Ragonnet-Cronin et al. 2013) or as a percentile of the distribution of pairwise sequence distances (Prosperi et al. 2011). A fundamental limitation of these non-parametric phylogenetic clustering tools is the need to define this arbitrary absolute transmission cluster divergence thresholds (termed as ‘cutpoints’ by Villandre et al., 2016). The lack of a consensus definition of a phylogenetic transmission cluster (Grabowski and Redd 2014) coupled with incomplete knowledge of a pathogen’s underlying epidemiological dynamics often reduces the choice of cutpoints to an *ad hoc* exploratory exercise resulting in subjective cluster definitions.

Phydelity is a novel phylogenetic clustering tool designed to negate the need for arbitrarily defined cluster divergence thresholds. Requiring only the phylogenetic tree as input, Phydelity infers putative transmission clusters through the identification of groups of sequences that are more closely-related to one another than the ensemble distribution under a statistically-principled framework. Phydelity, like another phylogenetic clustering tool that we recently developed, PhyCLIP, is based on integer linear programming (ILP) optimisation (Han et al. 2019). However, the two clustering tools are substantially different in their approaches and ILP models such that their clustering results have entirely distinct interpretations. PhyCLIP uses the divergence information of the entire phylogenetic tree to inclusively assign statistically-supported cluster membership to as many sequences in the tree as possible that putatively capture variant ecological, evolutionary or epidemiological processes. To this end, PhyCLIP is useful for sub-species nomenclature development. Phydelity, on the other hand, exclusively distinguishes closely-related pathogens with pairwise sequence divergence that are significantly more likely to be drawn from the same low divergence distribution than that of the ensemble. As such, while PhyCLIP’s designated clusters are underpowered to be interpreted as sequences linked by transmission events, clusters inferred by Phydelity can be interpreted as putative transmission clusters (see Supplementary Materials).

To demonstrate the utility of Phydelity in identifying putative transmission clusters, the algorithm underlying Phydelity is first presented in detail. The clustering tool is then applied to both simulated and empirical datasets, including outbreaks of Hepatitis B and C viruses as well as seasonal A/H3N2 influenza virus infections, and compared against results generated by existing phylogenetic clustering methods. Phydelity is freely available at http://github.com/alvinxhan/Phydelity.

## Method

### Clustering Algorithm

Figure 1(a) shows the overall workflow of Phydelity. First, Phydelity considers the input phylogeny as an ensemble of putative clusters, each consisting of an internal node *i* and the leaves it subtends. The within-cluster diversity of node *i* is measured by its mean pairwise patristic distance (μ_*i*_). The patristic distance between two nodes, which can be any sequence tips or internal nodes in the phylogeny, refers to the sum of branch lengths linking those two nodes. Sequences subtended by *i* (i.e. all descendant tree tips of node *i*) are considered for clustering if μ_*i*_ is less than the maximal patristic distance limit (*MPL*), under which sequences are considered more closely-related to one another than the ensemble distribution (Figure 1b).

**Figure 1.**
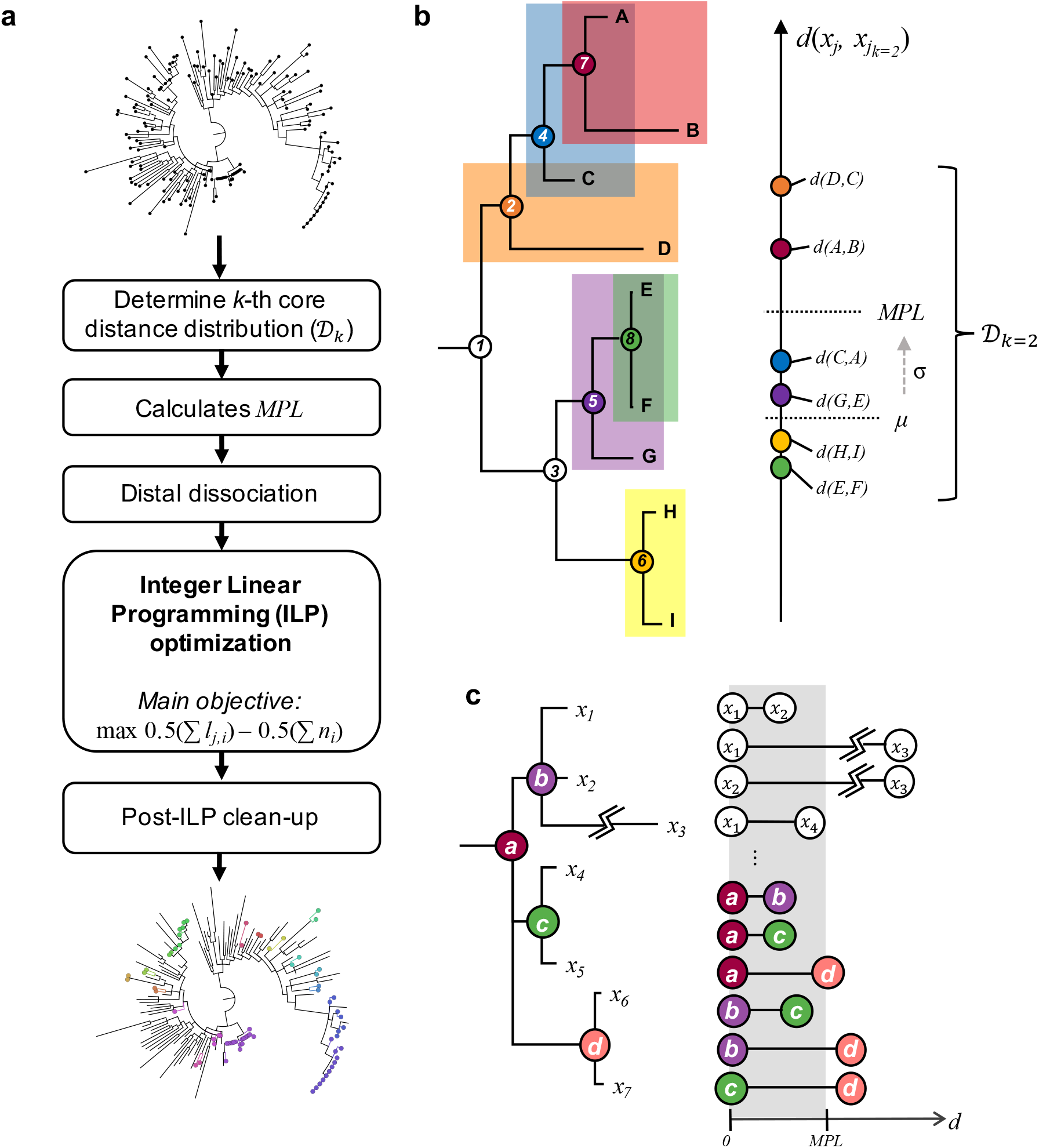
**(a)** Phydelity algorithm pipeline. Phydelity considers the input phylogenetic tree as a collection of putative clusters each defined by an internal node *i* and tips *j* that it subtends. The algorithm first infers the *k*-th core distance distribution (𝒟_*k*_) from the pairwise patristic distances of the closest *k*-neighbouring tips. *k* can be defined by the user or scaled by Phydelity to obtain the supremum 𝒟_*k*_ with the lowest divergence. 𝒟_*k*_ is then used to compute the maximal patristic distance limit (*MPL*) under which tips are considered to be more closely-related than to the ensemble. Dissociation of distally related subtrees/sequences (Figure 1c) ensues such that both monophyletic and paraphyletic clustering structures can be identified. Phydelity then incorporates the distance and topological information of the remaining nodes and tips into an integer linear programming (ILP) model to be optimised by clustering all tips that satisfy the relatedness constraints within the least number of clusters. Finally, post-ILP steps are implemented to remove any tips that may have been spuriously clustered. **(b)** Determination of the maximal patristic distance limit (*MPL*) using the median (μ) and robust estimator of scale (σ) based on the *k*-th core distance distribution (𝒟_*k*_) of every sequence *xj* and its *k*-closest neighbours 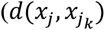; *k*=2 in this case as shown by the pairs of sequences highlighted with distinct colours). **(c)** Distal dissociation of a putative transmission cluster subtended by internal node ***a***. If a sequence tip has a pairwise sequence distance that is greater than *MPL*, it will be dissociated and not be clustered under the internal node of interest (i.e. internal node ***a***). In this case, sequence *x*_3_ is dissociated from the putative cluster ***a*** due to its exceedingly long branch length violating the *MPL* threshold (i.e. 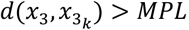). Additionally, whole subtrees subtended by the internal node of interest will be dissociated if any of its inter-nodal patristic distance exceeds *MPL*. Here, subtree ***d*** and its descending sequences (i.e. *x*_6_ and *x*_7_) will be dissociated from ***a*** as its inter-nodal distances with internal nodes ***b*** and ***c*** are both larger than *MPL*.

Phydelity computes the *MPL* by first calculating the pairwise patristic distance distribution of closely-related tips comprising the pairwise patristic distances of sequence *x*_*j*_ to the closest *k*-neighbouring tips (i.e. 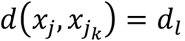) wherein their closest *k*-neighbours include sequence *x*_*j*_ as well (i.e. the *k*-th core distance distribution, 𝒟_*k*_; Figure 1b). Additionally, 𝒟_*k*_ is incrementally sorted (*d*_*l*_ ≤ *d*_*l+1*_) and truncated up to *d*_*L*_ if the common log difference between *d*_*L*_ and *d*_*L+1*_ is more than zero:

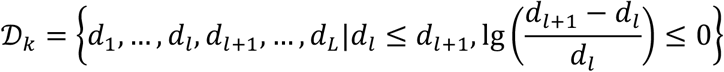

The user can opt to either input the desired *k* parameter or allow Phydelity to automatically scale *k* to the value that yields the supremum *k*-th core distance distribution with the lowest overall divergence (i.e. the largest possible *k* that still yields the lowest overall divergence between *k*-neighbouring tips). This is done by testing if 𝒟_*k+1*_ and 𝒟_*k*_ are statistically distinct (*p* < 0.01) using the Kuiper’s test (see Supplementary Materials). All clustering results of Phydelity presented in this work were generated using the autoscaled value of *k*.

The *MPL* is then calculated by:

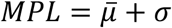

where 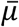 is the median pairwise distance of 𝒟_*k*_ and σ is the corresponding robust estimator of scale without assuming symmetry about 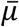 using the Qn method (see Supplementary Materials, Rousseeuw and Croux 1993; Figure 1b).

This is then followed by dissociation of distantly-related descendant subtrees/sequences to all putative nodes for clustering, thereby facilitating identification of both monophyletic as well as nested paraphyletic clusters (Figure 1c; see Supplementary Materials). Phydelity filters outlying tips from putative clusters under the assumption that viruses infecting individuals in a transmission chain coalesce to the same most recent common ancestor (MRCA).

Additionally, Phydelity requires any clonal ancestors in between the MRCA and tips of a putative cluster to be as genetically similar to each other as they are to the tips of the cluster. As such, for a putative transmission cluster, the mean pairwise nodal distance between all internal and tip nodes of a cluster must also be ≤ *MPL* (Figure 1c).

An ILP model is implemented and optimised under the objective to assign cluster membership to sequences satisfying the aforementioned relatedness criteria within the least number of clusters. In other words, Phydelity uses ILP optimisation to search for the clustering configuration that favours the designation of larger clusters of closely-related sequences which are likely linked by transmission events. Any topologically outlying singletons that were spuriously clustered are removed. Finally, it is important to note that a transmission cluster identified by Phydelity should only be interpreted as a fully connected network of likely transmission pairs without implying any underlying transmission directionality. The full algorithm description and mathematical formulation of Phydelity is detailed in the Supplementary Materials.

### Assessing clustering results of simulated epidemics

Phydelity was evaluated on phylogenetic trees derived from simulated HIV epidemics of a hypothetical men who have sex with men (MSM) sexual contact network (C-type networks in Villandre *et al.*, 2016). The simulated sexual contact network comprised 100 subnetworks (communities) sampled from an empirical distribution obtained from the Swiss HIV Cohort Study. All communities were linked in a chain initially and additional connections between any two communities were generated at a probability of 0.00075. Subjects in the network could either be in the “susceptible”, “infected” or “removed” (i.e. individual was diagnosed and sampled) state. Transmission clusters were attributed to sexual contact among individuals belonging to the same community.

300 epidemics were simulated for four different weights of inter-community transmission rates (*w* = 25%, 50%, 75% or 100% of the within-community rate). Two infected individuals were randomly introduced in any of the 100 communities. Transmission time along an edge followed an exponential distribution with rates directly proportional to the associated weights. Time until removal was based on a shifted exponential distribution with the shift representing the minimum amount of time required for a virus to be transmitted to susceptible neighbours. The simulation ended once 200 individuals were in the “removed” state.

These simulated datasets were tested by Villandre *et al.* (2016) to compare the outputs of four “cutpoint-based” phylogenetic clustering methods where the arbitrary distance threshold defining a transmission cluster (i.e. cutpoint) was computed as the: (i) absolute patristic distance threshold between any two tips (Brenner et al. 2007); (ii) standardised number of nucleotide changes (i.e. ClusterPicker, Ragonnet-Cronin *et al.*, 2013); (iii) percentile of the phylogeny’s pairwise sequence patristic distance distribution (i.e. PhyloPart, Prosperi *et al.*, 2011) and (iv) height of an ultrametric tree obtained using the weighted pair-group method of analysis (WPGMA). For each method, Villandre *et al*. varied the corresponding cutpoint parameter over an equivalent range of thresholds. Comparing the output clusters generated by the four methods at their respective optimal cutpoint by adjusted rand index (see below), it was found that the WPGMA method tended to produce clusters with better correspondence to the underlying sexual contact structure. As such, clustering results from Phydelity were compared to those obtained by Villandre *et al*. using the WPGMA method. Additionally, Phydelity was also compared to the multi-state birth-death (MSBD) method which inferred transmission clusters on the same simulated datasets by detecting significant changes in transmission rates (Barido-Sottani et al. 2018).

To assess and compare the output clusters from Phydelity and the aforementioned clustering methods that had been tested on these networks previously, several metrics were used to measure how well the clustering results corresponded with the known sexual contact network:

i. Adjusted rand index (*ARI*). ARI measures the accuracy of the clustering results by computing the frequencies of pairs of sequences of the identical (or distinct) subnetwork(s) assigned to the same (or different) cluster(s) (Hubert and Arabie 1985). ARI ranges between −1 (matching between output clusters and community labels is worse than random clustering) and 1 (perfect match between output clusters and ground truth).
ii. Modified Gini index (*I*_*G*_). Gini impurity, commonly used in decision tree learning, refers to the probability of a randomly selected item from a set of classes being incorrectly labelled if it was randomly labelled by the distribution of occurrences in the class set (Breiman et al. 1984). Here, *I*_*G*_ measures how often a randomly selected sequence from the given network would be incorrectly clustered by the inferred clusters. For a sexual contact network with *T* communities (i.e. *t* ∈ {1, 2, …, *T*}), *I*_*G*_ is computed as:

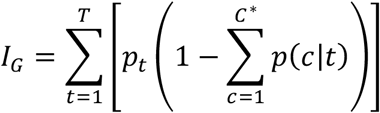

where *C*^∗^ is the set of clusters defined to have correctly classified sequences attributed to community *t* (i.e. any cluster that constitutes the largest proportion of sequences from community *t* at both the cluster and the community label levels), *p*_*t*_ is the probability of sequence from community *t* and *p*(*c*|*t*) refers to the probability that a sequence is clustered under cluster *c* conditional of it being from community *t*. If output clusters perfectly align with the underlying sexual contact network (i.e. one cluster only constitute one class of community), *I*_*G*_ = 0. Conversely, if clustering results are completely random, *I*_*G*_ = 1.
iii. Purity measures the average extent that the output clusters contain only a single class (i.e. a particular sexual contact community; Manning et al. 2008):

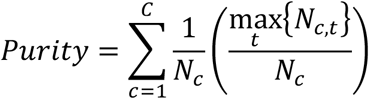

where *N*_*c*_ is the size of cluster *c*, *N*_*c,t*_ is the number of tips from community *t* clustered under cluster *c* and *C* is the set of all output clusters. Note that purity (as well as *I*_*G*_) can be inflated if the total number of clusters is large (i.e. if each tip is assigned to a unique cluster, purity = 1 and *I*_*G*_ = 0).
iv. Normalised mutual information (*NMI*) trades off the output clustering quality against the number of clusters (Manning et al. 2008):

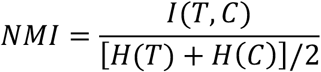

where *H(T)* and *H(C)* are the respective entropies of the network communities and output clusters, and *I(T,C)* is the mutual information between them. If clustering is random with respect to the network community labels, *I(T,C)* = 0 (i.e. *NMI* = 0). On the other hand, maximum mutual information is achieved (i.e. *I*(*T*, *C*) = *I*(*T*, *C*)_*max*_) either when the output clusters map the sexual contact network perfectly or all clusters have one member only. Hence, to penalise large cardinalities (i.e. number of members in a cluster) while normalising *I*(*T*, *C*) between 0 and 1, *NMI* is calculated since (a) entropy increases with increasing number of clusters and (b) [*H*(*T*) + *H*(*C*)]/2 is a tight upper bound to *I*(*T*, *C*).

### Empirical datasets

Phydelity was also tested on three empirical datasets – acute hepatitis C virus infections among men who have sex with men (Charre et al. 2018), hepatitis B viruses collected from members of the same families (Matsuo et al. 2017) as well as A/H3N2 influenza viruses collected from a community-based cohort of households during the 2014/2015 season (McCrone et al. 2018). All phylogenetic trees were reconstructed using RAxML (v8.2.12) under the GTRGAMMA model (Stamatakis 2014).

### Comparisons to ClusterPicker and PhyloPart

ClusterPicker (Ragonnet-Cronin et al. 2013) and PhyloPart (Prosperi et al. 2011), two non-parametric phylogenetic clustering tools that are methodologically comparable to Phydelity, were also applied to the hepatitis C and hepatitis B virus datasets for comparisons. Either clustering tool has been previously applied to multiple studies involving different pathogens (Prosperi et al. 2011; Jacka et al. 2014; Bezemer et al. 2015; Bartlett et al. 2016; Coll et al. 2017; de Oliveira et al. 2017; Charre et al. 2018). Other than the phylogenetic tree, both ClusterPicker and PhyloPart also require users to input an arbitrarily-defined genetic distance threshold (as an absolute distance limit for ClusterPicker and percentile of the global pairwise patristic distance for PhyloPart). As such, a range of distance limits (PhyloPart: 0.5-10^th^ percentile; ClusterPicker: 0.005-0.1 nucleotide/site) were applied to both tools. No bootstrap support threshold were implemented for comparability to Phydelity.

The lowest optimal threshold for the distance range tested was found by maximisation of the mean silhouette index (*SI*) for both ClusterPicker and PhyloPart. The Silhouette index measures how similar an item is to members of its own cluster as opposed to the nearest neighbouring clusters - i.e. a larger mean silhouette index indicates that items of the same cluster are more closely related amongst themselves than to its neighbours (Rousseeuw, 1987). No parameter optimisation was required for Phydelity.

## Results

### Simulated HIV epidemics

Phydelity was applied to simulated HIV epidemics among men who have sex with men (MSM) belonging to a hypothetical sexual contact network structures where transmission clusters were attributed to transmission by sexual contact among individuals belonging to the same subnetwork (see Methods; Villandre *et al.*, 2016). These simulations were originally used to assess the performance of “cutpoint-based” clustering tools, including ClusterPicker, PhyloPart as well as the weighted pair-group method of analysis (WPGMA) which generally attained the highest adjusted rand-index (ARI) score across all simulations when calibrating their respective cutpoint thresholds against the ground-truth. Phylogenetic trees generated from these simulations were also tested by the multi-state birth death (MSBD) method (Barido-Sottani et al. 2018).

Clustering results from Phydelity were compared to outputs from the MSBD method and those from the WPGMA method achieving the best ARI scores. The purity, modified Gini index (*I*_*G*_) and normalised mutual information (*NMI*) measures were also used to provide a more comprehensive assessment of the clustering results (Figure 2, Supplementary Figure 3 and Supplementary Table 1; see Methods).

**Figure 2.**
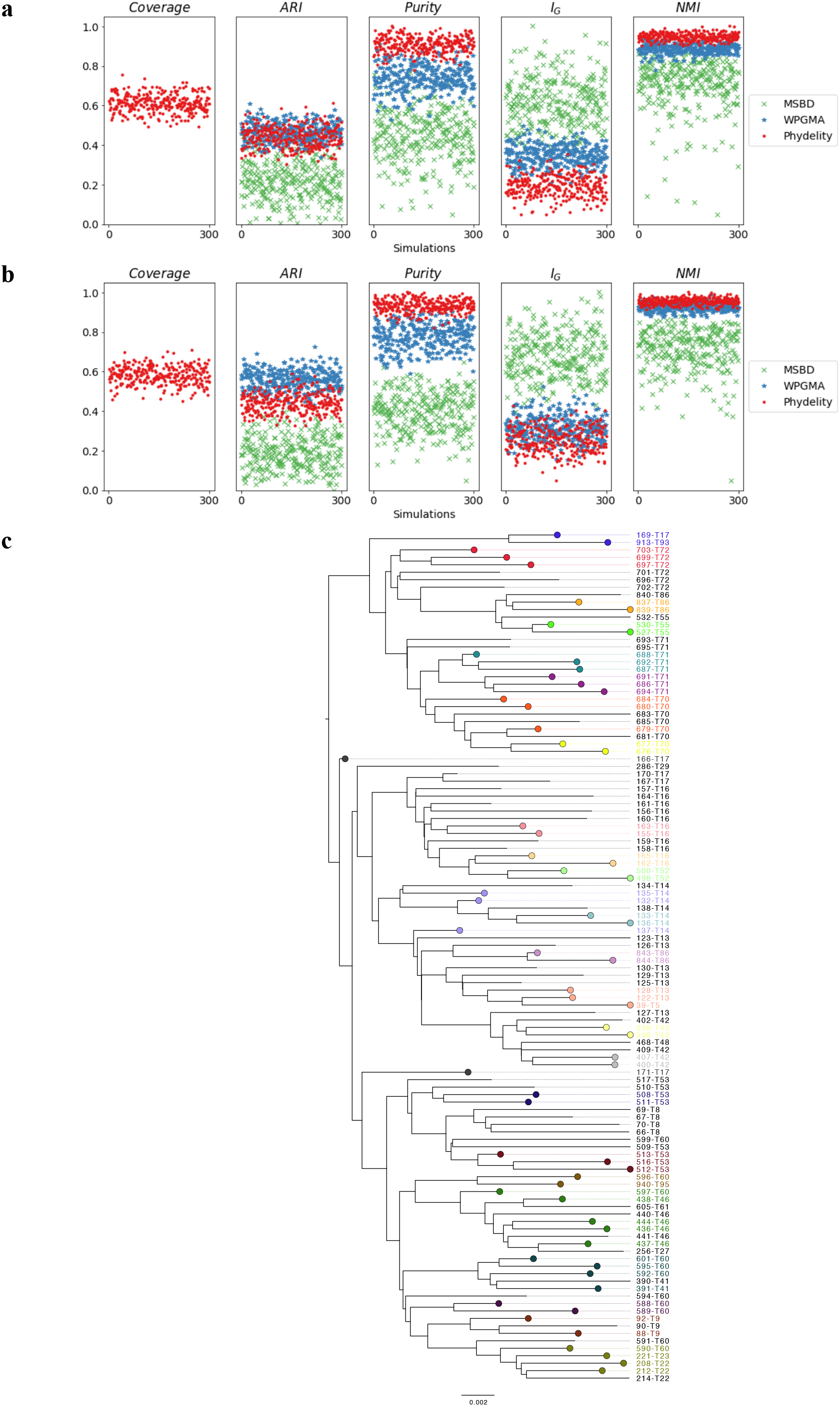
Clustering results of simulated HIV epidemics in a hypothetical MSM sexual contact network. **(a)** Clustering metrics for clustering algorithms (Phydelity, weighted pair-group method of analysis (WPGMA) and multi-state birth death (MSBD) methods) applied simulated phylogenies with inter-communities transmission rates weighted at half of within-community rates (i.e. *w* = 0.5). Coverage refers to the proportion of tips clustered by Phydelity. Adjusted rand index (*ARI*) measures how accurate the output clusters corresponded with the community labels. *Purity* gives the average extent clusters contain only a single class of community. Modified Gini index (*I*_*G*_) is the probbility that a randomly selected sequence would be incorrectly clustered. Normalised mutual information (*NMI*) accounts for the trade-off between clustering quality and number of clusters. **(b)** Results for simulations where inter-communities transmission rates were identical to within-community rates (i.e. *w* = 1.0). **(c)** Sample output clusters of Phydelity for a subtree of an example simulation (*w* = 0.5). Tips that were clustered by Phydelity are distinctly coloured according to their cluster membership. By relaxing the monophyletic assumption, Phydelity is capable of detecting paraphyletic clusters (e.g. transmission pair 166-T17 and 171-T17 and cluster subtending 132-T14, 135-T14 and 137-14).

The phylogenetic trees generated from the simulations had a large number of clusters that were relatively small in size (i.e. percentage of sequences that were part of ground truth clusters with sizes < 8 tips = 33.9% (weight of inter-community transmission rates*, w* = 25%); 55.5% (*w* = 100%); see Barido-Sottani *et al.* (2018) for more details). Furthermore, these ground truth clusters were not all monophyletic (Figure 2c). As a result, while Phydelity and WPGMA yielded comparable ARI scores (Phydelity: 0.44-0.45 (s.d. = 0.05); WPGMA: 0.44-0.56 (s.d. = 0.05-0.05); Supplementary Table 1), Phydelity’s output clusters, which allows paraphyletic clusters (Figure 2c), are substantially purer (mean purity; Phydelity: 0.81-0.88 (s.d. = 0.03); WPGMA: 0.67-0.74 (s.d. = 0.06-0.06)) and have a lower probability of misclassification when compared to WPGMA which assumes clusters are strictly monophyletic (mean *I*_*G*_; Phydelity: 0.27-0.28 (s.d. = 0.04-0.05); WPGMA: 0.33-0.40 (s.d. = 0.04-0.05)). Coverage of sequences clustered by Phydelity lies between 58.2% and 61.6%.

The clustering results from WPGMA presented in this work were based on the optimal distance threshold derived by calibration against the simulated ground-truth. Notably, Phydelity’s auto-scaling mitigates the need for threshold calibration and enables application to empirical datasets where ground truth clustering is unavailable, as is typically the case for epidemiological studies.

### Hepatitis B virus transmission between family members

Phydelity was tested on empirical datasets to demonstrate its applicability on real-world data, including hepatitis B viruses (HBV) collected from residents in the Binh Thuan Province of Vietnam (Matsuo et al. 2017). In such highly endemic regions, HBV is commonly transmitted either vertically from mothers to children during the perinatal period or horizontally between cohabitants of the same household (Matsuo et al. 2017). As complete genome nucleotide sequences were not available for all individuals, a phylogenetic tree was reconstructed using the viral polymerase sequences collected from 41 patients, of which 12 of them were confirmed to be members of three families (i.e. denoted as F2, F3 and F4) by a family survey as well as mitochondrial analyses. Besides Phydelity, the resulting phylogeny was also implemented in ClusterPicker and PhyloPart.

Phydelity identified three likely transmission clusters that distinguish between the separate family households (Figure 3). At their respective optimal distance thresholds by mean Silhouette index (see Methods), ClusterPicker and PhyloPart achieved similar clustering results. Importantly, Phydelity was able to obtain the same optimal clustering results without optimisation and implementation of a hard-to-interpret distance parameter.

**Figure 3.**
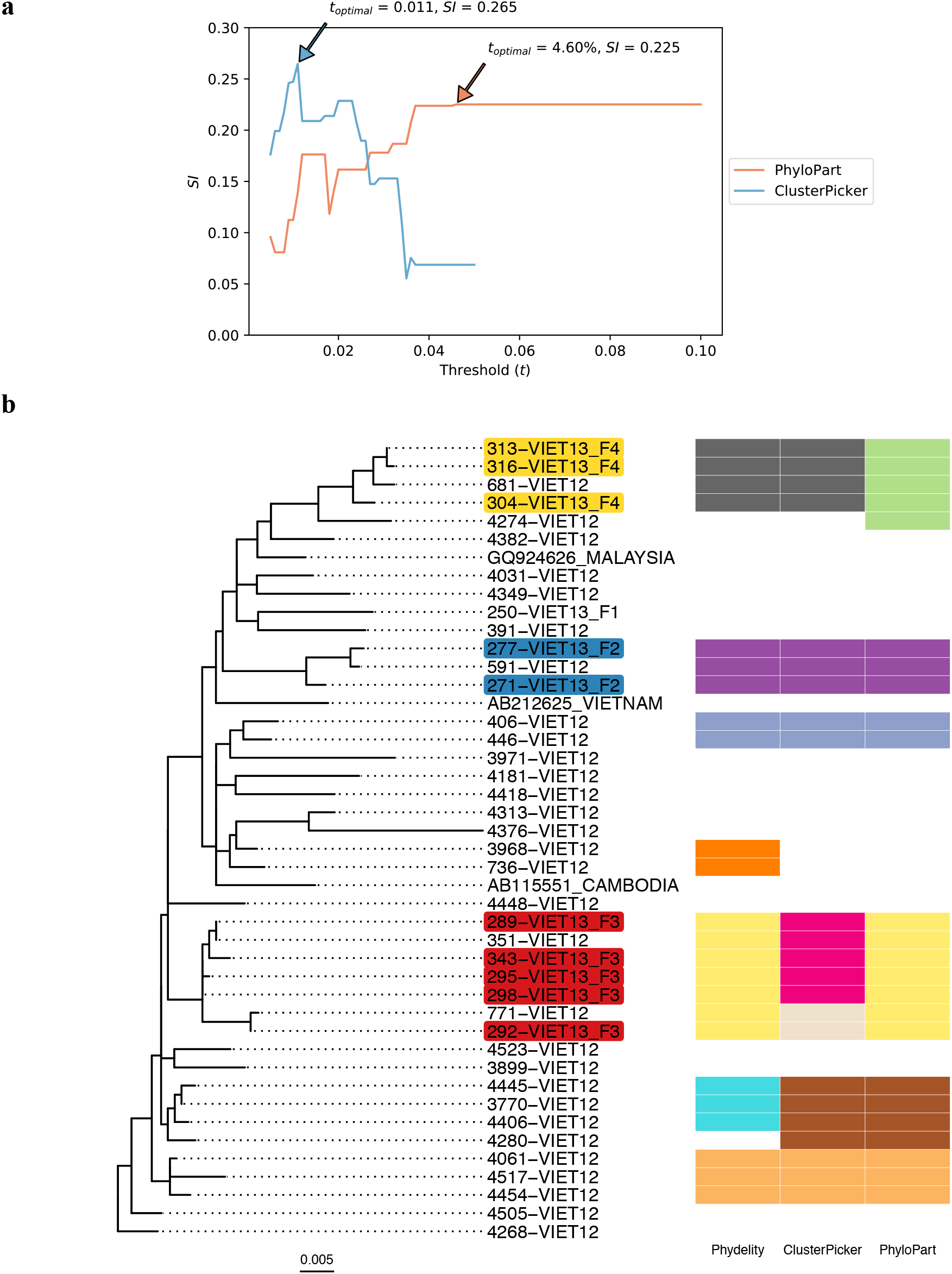
Clustering results of hepatitis B viruses (HBV) collected from residents in the Binh Thuan Province of Vietnam. (**a**) Plots of mean Silhouette index (*SI*) computed for the range of genetic distance thresholds implemented in ClusterPicker and PhyloPart. Clustering results from the lowest optimal distance threshold (*t*_*optimal*_) with the highest *SI* value for each method were compared to Phydelity as depicted in **b** (ClusterPicker: *t*_*optimal*_ = 0.011 nucleotide/site, *SI* = 0.265; PhyloPart: *t*_*optimal*_ = 4.60%, *SI* = 0.225). Plot for ClusterPicker is truncated at ~0.05 nucleotide/site as the entire tree collapsed to single cluster after this threshold. (**b**) Maximum likelihood phylogeny of HBV polymerase sequences derived from viruses collected from 41 patients. 12 patients were confirmed to be members of three separate family households (F2, F3 and F4; tip names shaded with a distinct colour for each family). Clustering results from Phydelity are depicted as a heatmap alongside outputs from ClusterPicker and PhyloPart based on their respective *t*_*optimal*_. Each distinct colour of the heatmap cells denotes a different cluster.

### Hepatitis C virus transmission among MSM

Incidence of HCV infections among HIV-negative MSM has been relatively limited as compared to their HIV-positive counterparts. However, the recent uptake of pre-exposure prophylaxis (PrEP) among HIV-negative individuals to prevent HIV infection could pose higher risk of sexually transmitted HCV infections (Volk et al. 2015; Charre et al. 2018). In a study on HIV-positive and HIV-negative MSM patients in Lyon, 108 cases of acute HCV infections (80 primary infections; 28 reinfections) were reported between 2014 and 2017 among 96 MSM (72 HIV-positive; 24 HIV-negative, of which 16 (67%) of them were on PrEP; Charre et al. 2018). Separate phylogenetic analyses were performed on a subset of 89 (68 HIV-positive; 21 HIV-negative) HCV isolates belonging to genotypes 1a and 4d based on their NS5B sequences. Additionally, 25 HCV sequences from HIV-infected MSM collected before 2014 were included along with 60 control HCV sequences derived from HIV-negative, non-MSM patients residing in the same geographical area as controls. All sequences collected from MSM patients were given strain names in the format of “MAH(ID)_accession” while control sequences from non-HIV, non-MSM patients were denoted as “NCH(ID)_accession” (Figure 4). Phydelity as well as ClusterPicker and PhyloPart were applied to the reconstructed phylogenies, with the latter calibrated over a range of distance thresholds. Again, only clustering results based on the lowest distance threshold maximising the mean Silhouette index for ClusterPicker and PhyloPart were compared to Phydelity’s output clusters (see Methods).

**Figure 4.**
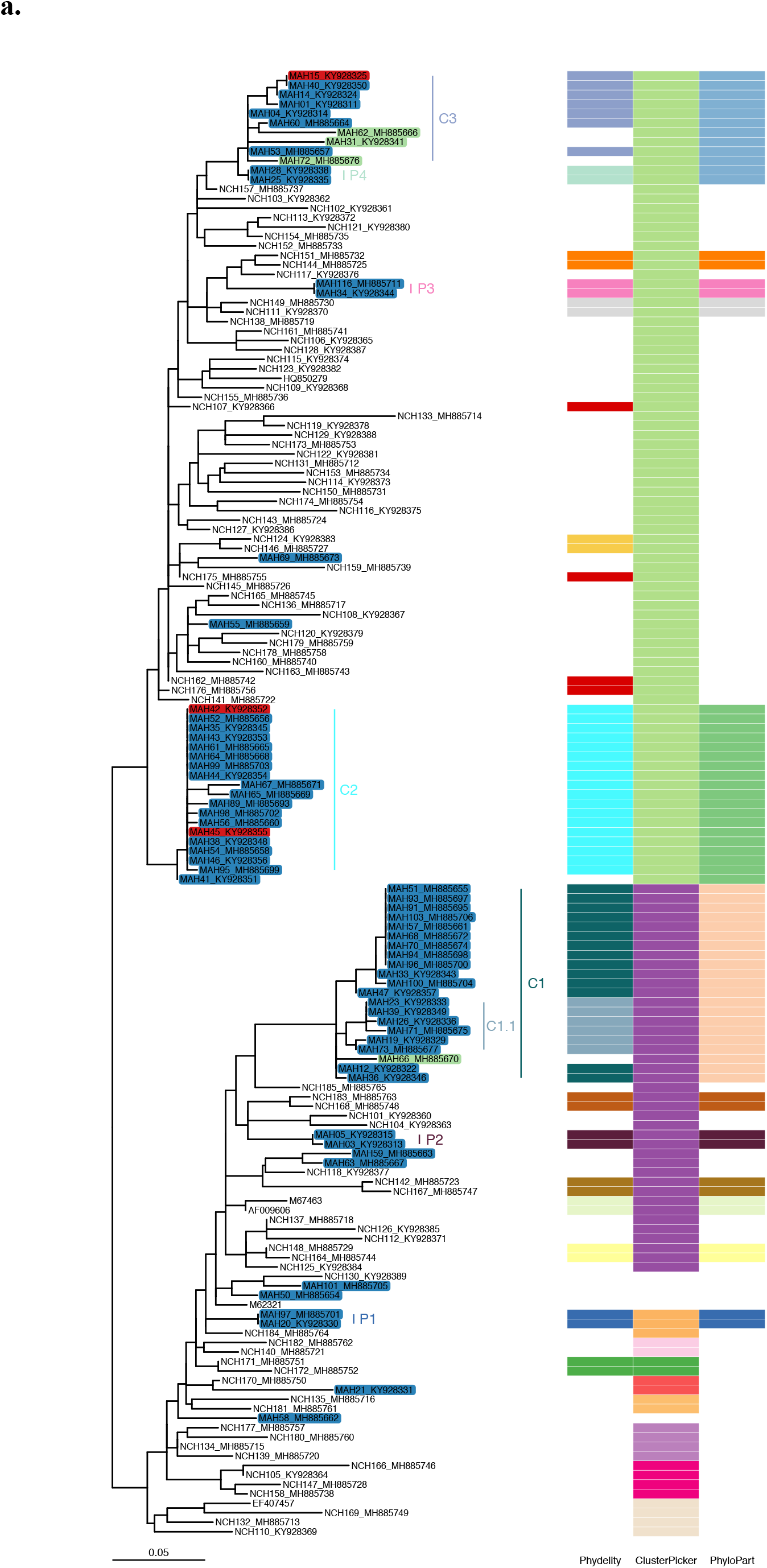
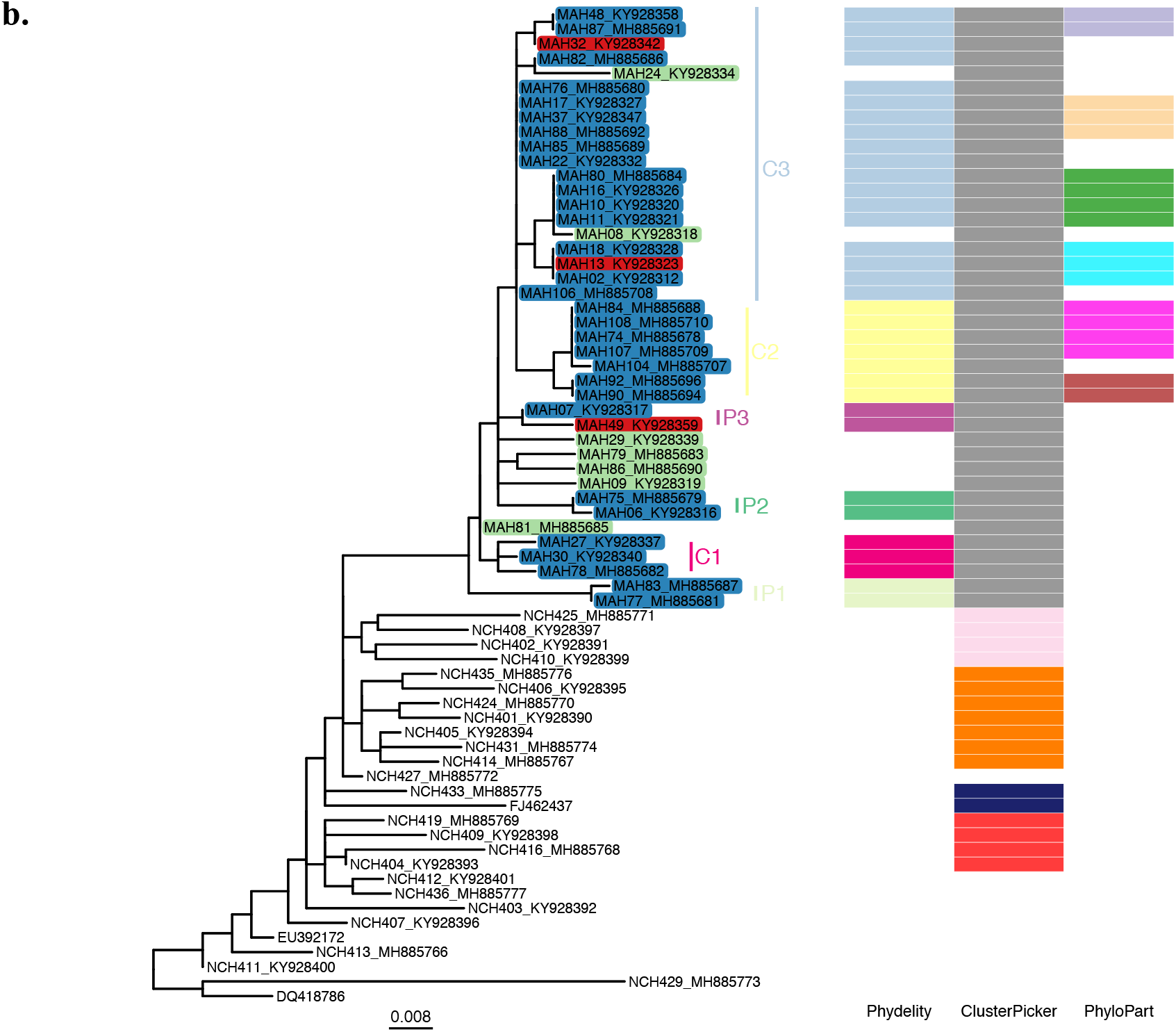
Maximum likelihood phylogeny and clustering results of hepatitis C viruses (HCV) obtained from men who have sex with men (MSM) in Lyon, France. All highlighted tip names denoted in the format “MAH(ID)_accession” were samples from MSM patients (blue: HIV-positive, red: HIV-negative, green: HIV-positive and considered as outlying sequences by Phydelity). Non-highlighted tips were collected from non-HIV, non-MSM patients residing in the same geographic region and time period. Clustering results from Phydelity, ClusterPicker and PhyloPart are depicted as a heatmap. Each distinct colour refers to a different cluster. Similar to the Vietnamese hepatitis B empirical viral datasets (Figure 3a and Supplementary Figure 4), mean Silhouette index was used as the optimality criterion to determine the optimal absolute distance threshold for ClusterPicker and PhyloPart. Only results based on the optimised thresholds are shown here for ClusterPicker and PhyloPart. No parameter optimisation is required for Phydelity. **(a)** Genotype 1a. **(b)** Genotype 4d.

Generally, membership of the MSM transmission clusters and pairs identified by Phydelity across both genotypes were strictly limited to sequences derived from MSM patients. Relaxing the monophyletic assumption by dissociating distantly-related tips from putative monophyletic clusters (see Methods) enables Phydelity to identify likely outlying sequences as evidenced by their relatively longer branch lengths from the cluster ensemble (Table 1 and Figure 4; Genotype 1a: cluster C1 – MAH66 and cluster C3 – MAH31, MAH62 and MAH72; Genotype 4d: cluster C3 – MAH24 and MAH08). In particular, for genotype 1a, even though the mean pairwise distance of MAH72 to members of cluster C3 is within a standard deviation of the latter’s within-cluster diversity, its distance to the more distant members (e.g. MAH15 and MAH40, Figure 4) violated the inferred *MPL* (Table 1). Additionally, as a result of distal dissociation, Phydelity distinguishes clusters that are genetically more alike amongst themselves than to those phylogenetically ancestral to it (e.g. cluster C1.1 that is “nested” within cluster C1 for genotype 1a; Figure 4a).

**Table 1:** Comparing the genetic distance between outlying tips and the clusters they coalescence to with the genetic diversity of those clusters.

| Genotype | MPL | Cluster | Mean pairwise patristic distance of cluster ($\sigma$) | Outlier | Mean pairwise patristic distance of outliers to cluster members ($\sigma$) |
| --- | --- | --- | --- | --- | --- |
| 1a | 0.029 | C1 | 0.011 (0.012) | MAH66 | 0.043 (0.009) |
|  |  | C3 | 0.016 (0.009) | MAH62 | 0.045 (0.027) |
|  |  |  |  | MAH31 | 0.041 (0.025) |
|  |  |  |  | MAH72 | 0.022 (0.015) |
| 4d | 0.010 | C1 | 0.006 (0.004) | MAH24 | 0.019 (0.006) |
|  |  |  |  | MAH08 | 0.009 (0.005) |

For both genotypes, Phydelity found multiple clusters that included both HIV-positive and HIV-negative MSM patients (i.e. Genotype 1a: clusters C2 and C3, Figure 4a; Genotype 4d: clusters C2 and C2.2, as well as pair P2, Figure 4b). While it is not clear which of the HIV-negative patients were on PrEP (information not supplied in the original paper), the clustering results from Phydelity were in line with the findings by Charre *et al*. that acute HCV infections among HIV-negative MSM were likely sourced from their HIV-positive counterparts.

While ClusterPicker managed to consolidate all of the MSM genotype 4d sequences into a single monophyletic cluster (Figure 4b), its clustering of genotype 1a was problematic as a large number of non-MSM control sequences were clustered together with those from MSM patients (Figure 4a). PhyloPart’s optimal clustering output was consistent Phydelity’s for genotype 1a. However, the larger number of identical sequences in the genotype 4d tree skewed the optimal distance parameter (expressed as *x*-th percentile of the pairwise patristic distribution of the entire phylogeny) to only cluster these identical sequences.

### Seasonal A/H3N2 influenza virus infections within a community and the effects of sampling

Phydelity was also applied to A/H3N2 influenza viruses collected from a community-based cohort of 340 households (1431 participants) in Southeastern Michigan, U.S. during the 2014/2015 season (McCrone et al. 2018). Of the influenza positive cases, 206 virus samples were collected from 166 individuals that belonged to 81 households and sequenced. As concurrent infections among individuals within the same household do not necessarily imply transmission, McCrone et al. implemented stringent epidemiological as well as genetic distance constraints to identify transmission pairs: (i) the donor and recipient of a transmission pair were of the same household with onset of illness symptoms occurring within 7 days of each other, with the donor having the earlier symptom onset date; (ii) there must be no other potential donors with the same symptom onset date; (iii) symptom onset dates of donor and recipient should not be on the same day unless they were index cases; and (iv) genetic distance between the within-host viral populations of donor and recipient must be below the 5^th^ percentile of the distance distribution of random pairs of infected individuals from the community (McCrone et al. 2018). In total, 50 virus isolates constituting 32 high-quality transmission pairs were identified. Consolidating transmission pairs with overlapping donors and recipients into clusters, there were 22 genetically-validated transmission clusters, comprising of 16 pairs and 6 trios in total.

Using the phylogeny constructed from the consensus whole genome sequences of all 206 viruses, Phydelity was able identify 20 of the 22 high-quality transmission clusters as distinct clusters (Supplementary Figure 5). Applying the same metrics used to assess clustering performance of the simulated dataset earlier and using the high-quality transmission cluster labels as ground truth, Phydelity was able to produce highly pure clusters (97.8%), with a low probability of misclassification (*I*_*G*_ = 0.022) and good accuracy (*ARI* = 0.962), even after accounting for the number of predicted clusters (*NMI* = 0.993; Table 2). As transmission events defined by McCrone et al. were based on highly conservative criteria imposed on deep sequencing datasets, Phydelity, which operates at the consensus sequence level, could also cluster viruses that did not satisfy these constraints but were still linked epidemiologically by their household identities. As such, Phydelity’s clustering results were assessed based on the household association of the clustered individuals as well, yielding slightly diminished but nonetheless high quality performance (Purity = 0.894, *I*_*G*_ = 0.081, *ARI* = 0.791, *NMI* = 0.964; Supplementary Figure 5 and Table 2).

**Table 2:**
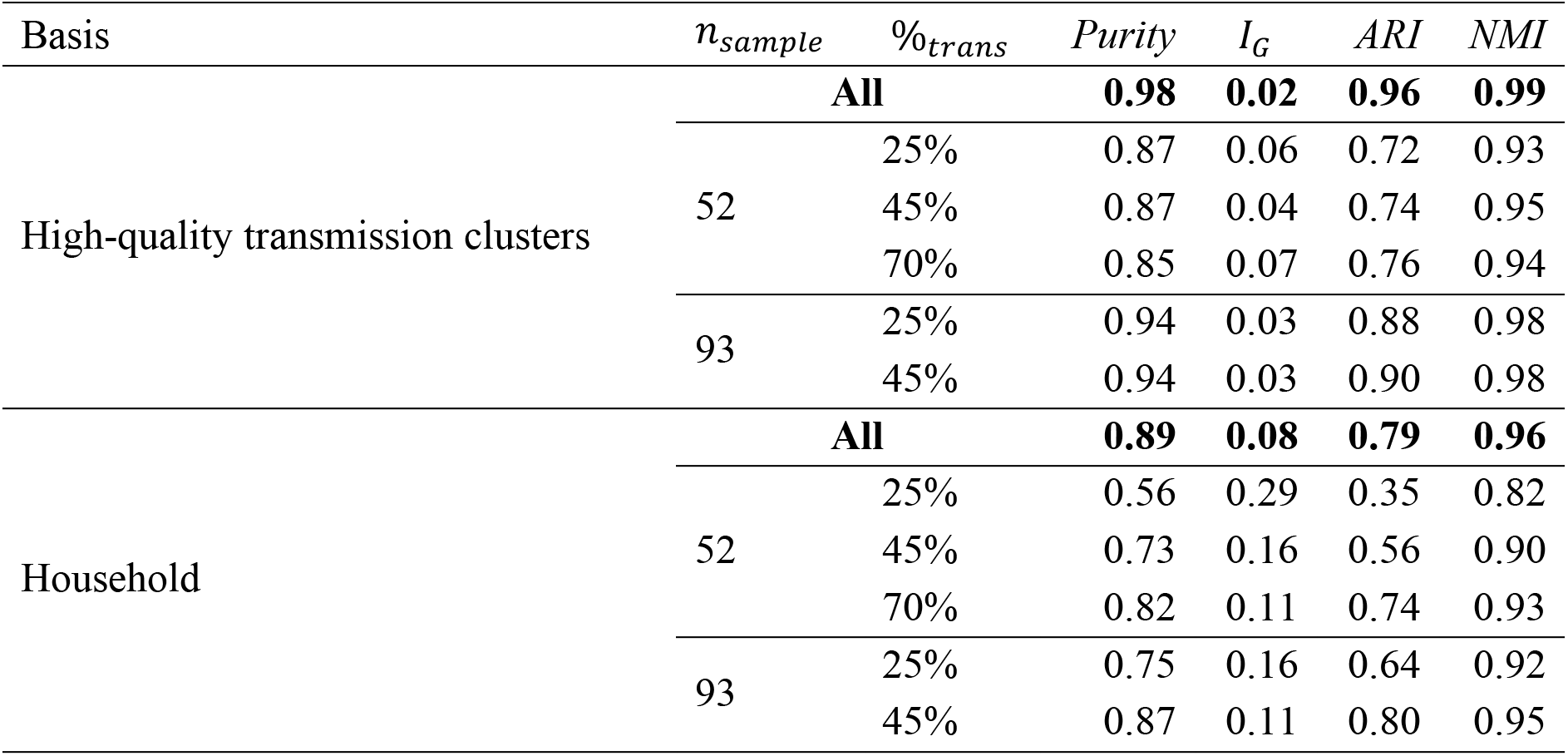
Clustering performance of Phydelity on seasonal A/H3N2 influenza viruses collected by McCrone et al. (2018). Ground truth used for clustering assessment is either based on the identities of genetically-validated, high-quality transmission clusters as defined by McCrone et al. or by the patients’ households. Besides analysing all of the viruses collected (bolded results), Phydelity was also applied to downsampled datasets consisting of different sample size (*n*_*sample*_) and proportion of sequences derived from the aforementioned high-quality transmission pairs (%_*trans*_). Adjusted rand index (*ARI*) measures how accurate the output clusters corresponded with the ground truth labels. *Purity* gives the average extent clusters contain only a single class. Modified Gini index (*I*_*G*_) is the probbility that a randomly selected sequence would be incorrectly clustered. Normalised mutual information (*NMI*) accounts for the trade-off between clustering quality and number of clusters.

| Basis | $n_{sample}$ | $\%_{trans}$ | *Purity* | $I_G$ | *ARI* | *NMI* |
| --- | --- | --- | --- | --- | --- | --- |
| High-quality transmission clusters | **All** |  | **0.98** | **0.02** | **0.96** | **0.99** |
|  | 52 | 25% | 0.87 | 0.06 | 0.72 | 0.93 |
|  |  | 45% | 0.87 | 0.04 | 0.74 | 0.95 |
|  |  | 70% | 0.85 | 0.07 | 0.76 | 0.94 |
|  | 93 | 25% | 0.94 | 0.03 | 0.88 | 0.98 |
|  |  | 45% | 0.94 | 0.03 | 0.90 | 0.98 |
| Household | **All** |  | **0.89** | **0.08** | **0.79** | **0.96** |
|  | 52 | 25% | 0.56 | 0.29 | 0.35 | 0.82 |
|  |  | 45% | 0.73 | 0.16 | 0.56 | 0.90 |
|  |  | 70% | 0.82 | 0.11 | 0.74 | 0.93 |
|  | 93 | 25% | 0.75 | 0.16 | 0.64 | 0.92 |
|  |  | 45% | 0.87 | 0.11 | 0.80 | 0.95 |

The full A/H3N2 sequence dataset was then randomly sampled to smaller pools of 52 (25%) as well as 93 (45%) isolates to assess how low sampling rates might affect Phydelity’s performance. To ensure that sequences involved in high-quality transmission pairs were also sampled, such isolates would constitute different proportions (either 25% or 45%; as well as 70% for pools of 52 sequences only) of the downsampled datasets. 10 distinct downsamples were generated for each sample size/high-quality transmission sequence combination and the average results were tabulated (Table 2).

As the *MPL* is informed by the phylogenetic tree, clustering results will consequently be sensitive to the diversity of closely-related tips within the input phylogeny. Specifically, the closely-related sequences that constitute the *k*-th core patristic distance distribution (𝒟_*k*_) must be homogenous (i.e. similar difference between consecutive distances when 𝒟_*k*_ is sorted; see Methods) but sufficiently distinct from the background diversity of the phylogeny. This was demonstrated by the improved clustering results with respect to household identities with greater proportional inclusion of genetically similar, high-quality transmission pairs in the downsampled dataset (Table 2). Furthermore, erroneous clustering of distantly-related tips can be obtained if 𝒟_*k*_ has a similar distance distribution relative to the entire tree due to insufficient divergence information from reduced sampling. This is evident from the general decrease in the clustering performance of all downsampled data. In particular, clustering closely-related, high-quality transmission clusters was worse off with a lower sample size.

### Computational performance

For computational performance, Phydelity can process a phylogeny of 1000 tips, on an Ubuntu 16.04 LTS operating system with an Intel Core i7-4790 3.60 GHz CPU, in ~3 minutes using a single CPU core and 253 MB of peak memory usage.

## Discussion

Phydelity is a statistically-principled tool capable of identifying putative transmission clusters from pathogen phylogenies without the need to introduce arbitrary distance thresholds. Instead, Phydelity infers the maximal patristic distance limit (*MPL*) for cluster designation using the pairwise patristic distance distribution of closely-related tips in the input phylogenetic tree. Unlike other cutpoint-based methods, Phydelity does not assume clusters are strictly monophyletic and can identify paraphyletic clustering owing to its distal dissociation approach. For datasets that span extended periods of time, multiple introductions within the same contact network and concurrent onward transmissions to other communities can result in “nested” introduction events that would go undetected by monophyletic clustering (Barido-Sottani et al. 2018). By relaxing this assumption, not only can Phydelity pick up these “nested” events, it tends to produce clusters that are purer with a lower chance of misclassification while excluding putative outlying tips that are exceedingly distant from the inferred cluster.

Even though there are algorithmic similarities between PhyCLIP (Han et al. 2019) and Phydelity, clustering results generated by PhyCLIP should not be interpreted as sequences linked by transmission events. For instance, when applied to the HCV genotype 1a NS5B dataset, PhyCLIP clustered 131 of the 155 input sequences into seven clades, all of which encompasses genetically similar viruses of both MSM and non-MSM origins that were endemic in Lyon during a specific period in time. In contrast, Phydelity assigned 73 sequences into 12 transmission pairs and 5 transmission clusters that distinguished the underlying MSM transmission events from non-MSM ones (Supplementary Figure 1). A detailed comparison between Phydelity and PhyCLIP can be found in Supplementary Materials.

One of the key assumptions made by Phydelity is that the transmitted pathogens coalesce to the same most recent common ancestor (MRCA) and that the pairwise genetic distance of internal nodes found between the MRCA and the tips of the cluster to be bounded below *MPL*. While Phydelity does not explicitly equate the inferred phylogeny to a transmission tree, imposing a distance threshold between the internal nodes within a phylogenetic cluster may be construed as an implicit assumption that the internal nodes are representative of transmission events. There are important differences in the interpretation of phylogenetic and transmission trees. The former depicts the shared ancestry between the sampled tips while the latter represents the true transmission history between the transmitted pathogens (Pybus and Rambaut 2009; Ypma et al. 2013). It should be noted that Phydelity neither attributes any interpretation of transmission events to the internal nodes nor does it relate branch lengths of the phylogenetic tree, which correlate with the timing of coalescence, to transmission times. Restricting the distances between internal nodes below the *MPL* is strictly meant to increase conservatism in identifying clusters that are as closely-related as possible.

There have also been criticisms that non-parametric cluster identification by genetic similarity is biased towards the detection of recent infections as opposed to discerning variations in transmission rates between different subpopulations, which can be further exacerbated by oversampling (Poon 2016; Dearlove et al. 2017; Le Vu et al. 2018). While this caveat limits the interpretation of the inferred transmission clusters, it does not render all phylogenetic clustering tools obsolete. Phylogenetic clustering tools supplemented by epidemiological meta-data can still be used to systematically identify infection trends, potential risk factors and target subpopulations, as demonstrated by multiple epidemiological studies of different measurably-evolving pathogens (Matsuo et al. 2017; de Oliveira et al. 2017; Charre et al. 2018).

Additionally, constructing a phylogenetic tree can be a computational bottleneck for large sequence datasets. As an alternative, genetic distance-based clustering algorithms such as HIV-TRACE (Kosakovsky Pond et al. 2018) which negate the need to build a phylogenetic tree have becoming increasingly popular. However, HIV-TRACE still requires users to specify an arbitrary absolute distance threshold. Additionally, while it performed better than other existing phylogenetic clustering methods, HIV-TRACE did not preclude problems with bias towards higher sampling rates (Poon 2016).

Despite its limitations, clustering results generated by Phydelity for the simulation and empirical datasets in this study demonstrate its superior performance over current widely used phylogenetic clustering methods. Importantly, Phydelity obviates the need for users to define or optimise non-biologically-informed distance thresholds. Phydelity is fast, generalisable, and freely available at https://github.com/alvinxhan/Phydelity.

## Acknowledgements

We would like to thank Frits Scholer for his assistance with optimising Phydelity, as well as Jelle Koopsen and Velislava Petrova for their intellectual contributions.

## Data availability

Phydelity is freely available on https://github.com/alvinxhan/Phydelity. All simulated datasets were downloaded from Villandre et al. (2016). Genbank accession numbers of HBV polymerase sequences: AB212625, GQ924626, AB115551, LC57377-LC57378, LC60789-LC60790, LC63767, LC64366-LC64378, LC64380-LC64381, LC80779-LC80783, LC80785, LC80787-LC80800, and LC80802-LC80804. Genbank accession numbers of HCV NS5B sequences: AF9606, EF407457, HQ850279, EU392172, FJ462437, DQ418786, M62321, MH885654-MH885777, and KY928311-KY928401. The A/H3N2 influenza virus consensus sequences were downloaded from https://github.com/lauringlab/Host_level_IAV_evolution. Jupyter notebooks used to analyse both simulated and empirical datasets can be downloaded from https://github.com/alvinxhan/Phydelity/tree/master/manuscript.

## Funding

A.X.H. was supported by the A*STAR Graduate Scholarship programme from A*STAR to carry out his PhD work via collaboration between Bioinformatics Institute (A*STAR) and NUS Graduate School for Integrative Sciences and Engineering from the National University of Singapore. E.P. was funded by the Gates Cambridge Trust (Grant number: OPP1144). S.M.S. was supported by the A*STAR HEIDI programme (Grant number: H1699f0013) and Bioinformatics Institute (A*STAR).

## Notes

#### Summary of Updates

Expanded Introduction; Additional analyses using A/H3N2 seasonal influenza virus sequence data from McCrone et al. (2018)

https://github.com/alvinxhan/Phydelity/

