## Supplementary Materials for "Inferring putative transmission clusters with Phydelity"

### Algorithm and mathematical formulation

Phydelity considers the input phylogenetic tree as an ensemble of $N$ monophyletic putative clusters, each defined by an internal node $i$ subtending a set of sequences $L_i$. The within-cluster divergence of putative cluster $i$ is measured by its mean pairwise sequence patristic distance ($\mu_i$).

Phydelity then determines the $k$-th core pairwise patristic distance distribution ($\mathcal{D}_k$) of closely-related tips from the input phylogeny. This is done by parsing for the $k$-closest sequences (i.e. $\{x_{j_k}\}$) to each tip $x_j$, ordering them by their patristic distance $d(x_j, x_{j_k})$. If the $k$-neighbours of $\{x_{j_k}\}$ also includes $x_j$ itself, then $d\left(x_j, x_{j_{k'}}\right)$ where $k' \leq k$ will be included in $\mathcal{D}_k$.

$\mathcal{D}_k$ is then incrementally sorted (i.e. let $d_l = d(x_j, x_{j_k})$, then $d_l \leq d_{l+1}$) and truncated up to $d_L$ if the common log difference between $d_L$ and $d_{L+1}$ is more than zero:

$$\mathcal{D}_k = \left\{d_1, \dots, d_l, d_{l+1}, \dots, d_L \middle| d_l \leq d_{l+1}, \lg\left(\frac{d_{l+1} - d_l}{d_l}\right) \leq 0\right\}$$

Users can opt to input their desired $k$ parameter or allow Phydelity to scale $k$ automatically, yielding the supremum distribution of $\mathcal{D}_k$ with the lowest overall divergence. This is achieved by iteratively testing if $\mathcal{D}_{k+1}$ and $\mathcal{D}_k$ are statistically distinct from each other ($p < 0.01$) by the Kuiper's test. Unlike the more widely-used Kolmogorov Smirnov test to test for equality between one-dimensional continuous distributions, the Kuiper's test statistic, which is also non-parametric, incorporates both the greatest positive and negative deviations between the distributions tested. The Kolmogorov Smirnov test statistic is only defined by their maximum difference. As such, the Kuiper's test has greater sensitivity as compared to the Kolmogorov-Smirnov test as it is able to detect shifts to both the tails as well as the median of the distributions.

The maximal patristic distance limit (*MPL*) is then calculated by:

$$MPL = \tilde{\mu} + \sigma \qquad (1)$$

$\tilde{\mu}$ is the median pairwise distance of $\mathcal{D}_k$ and $\sigma$ is the corresponding robust estimator of scale without assuming symmetry about $\tilde{\mu}$ using the Qn method (Rousseeuw and Croux 1993). Qn is an alternative estimator of scale such as the commonly used median absolute deviation (MAD). Like MAD, Qn also has a breakdown point of 50% - the maximum tolerance to outlying data in the distribution before the estimator yields unreliable results. However, it is computed from the pairwise differences of all values in the given distribution and does not require a location estimate (i.e. the median, which is used in calculating MAD). It has been proven to be statistically more efficient when applied to both Gaussian and non-Gaussian distributions relative to the MAD (Rousseeuw and Croux 1993).

To facilitate identification of both paraphyletic and monophyletic clusters, Phydelity dissociates distal subtrees and sequences from any ancestral node $i$ where $\mu_i > MPL$. Starting from the most distant sequence to $i$, Phydelity first applies a leave-one-out strategy dissociating sequences, and the whole descendant subtree if every sequence subtended by it was dissociated, until the recalculated $\mu_i$ without the distal sequences falls below $MPL$. $\mu_i$ is recalculated for any node with changes to its sequence membership $L_i$ after dissociating these distantly related sequences.

Phydelity filters outlying tips from putative clusters under the assumptions that viruses infecting individuals in a rapid transmission chain likely coalesce to the same most recent common ancestor (MRCA). Additionally, Phydelity requires any clonal ancestors in between the MRCA and tips of a putative cluster to be as genetically similar to each other as it is between the tips of the cluster. As such, for a putative transmission cluster, the mean pairwise nodal distance between all internal and tip nodes of a cluster must also fall below $MPL$.

Phydelity then formulates the phylogenetic clustering problem as an integer linear programming (ILP) model. Let $n_1, n_2, \dots, n_i, \dots, n_N$ be the set of binary variables indicating if putative cluster $i$ satisfies the conditions for clustering ($n_i = 1$ if it clusters; $n_i = 0$ if otherwise). Each sequence $j$ subtended by putative cluster $i$ is also assigned a binary variable $l_{j,i}$ indicating if the sequence is clustered under $i$ ($l_{j,i} = 1$ if $j$ is clustered under node $i$; $l_{j,i} = 0$ if otherwise).

The ILP model is optimised to cluster all sequences that satisfy the distance bounds defining a putative transmission cluster to as little number of clusters as possible. This is achieved by solving for the following equal-weighted multi-objective:

$$\max \sum_{j,i} l_{j,i} - \sum_{i} n_i \tag{2}$$

subject to the following constraints:

$$l_{j,i} \leq n_i \qquad \forall\ j \in L_i, i \qquad (3)$$

Constraint (3) stipulates that sequence $j$ can be clustered under node $i$ if and only if $i$ is a potential cluster ($n_i = 1$).

$$l_{j,i} \leq 2 - n_i - n_k \qquad \forall\ j \in \{L_i, L_k\}, k; i > k \qquad (4)$$

If sequence $j$ is subtended by putative clusters $i$ and $k$, wherein $i$ descended from $k$ and both nodes are potential clusters ($n_i = n_k = 1$), constraints (3) and (4) stipulate sequence $j$ will not be clustered under the descendant node $i$. Implementing these constraints across all pairwise combinations of subtrees subtending sequence $j$ in turn constrains $j$ to be clustered under the most ancestral node $k$ possible.

$$\sum_i l_{j,i} \leq 1 \qquad \forall\ j \qquad (5)$$

Constraint (5) stipulates that each sequence can only be clustered under a single cluster, hence abrogating any fuzzy clustering.

$$C(n_i - 1) \leq \sum_j l_{j,i} - 2 \qquad \forall\ i \qquad (6)$$

where $C$ is any arbitrarily large positive constant. Constraint (6) requires all clusters to subtend at least a pair of sequences.

$$C(n_i - 1) \leq MPL - \mu_i \qquad \forall\ i \qquad (7)$$

Constraint (7) ensures that $\mu_i$ of all clusters fall below the stipulated $MPL$ limit.

### Algorithmic differences between Phydelity and PhyCLIP

As mentioned in the main text, both Phydelity and PhyCLIP (Han et al. 2019) are statistically-principled phylogenetic clustering algorithms based on integer linear programming optimisation. However, there are substantial algorithmic differences between the two such that the clusters recovered have distinctly different interpretations (Supplementary Figure 1).

PhyCLIP was developed to identify statistically-supported subpopulations in pathogen phylogenies that putatively capture variant ecological, evolutionary or epidemiological processes that could underlie sub-species nomenclature development. PhyCLIP, like Phydelity, operates on the pairwise patristic distance distribution of an input phylogeny. However, PhyCLIP informs its within-cluster divergence limit using the *global* distribution of patristic distances, rather than the *local* distribution of *k*-neighbouring tips as implemented in Phydelity (Supplementary Figure 2). PhyCLIP's use of the pairwise patristic distance distribution of the entire tree allows it to test against the background genetic diversity of the global population included in the phylogeny, indicating whether the putative clustered sequences are sufficiently more related to one another than to the rest of the dataset to be designated a distinct cluster. PhyCLIP calculates the within-cluster divergence limit ($WCL$) as:

$$WCL = \tilde{\mu} + (\gamma\sigma) \tag{8}$$

where $\tilde{\mu}$ is the grand median of the mean pairwise patristic distance distribution $\{\mu_1, \mu_2, \ldots, \mu_i, \ldots, \mu_N\}$ and $\sigma$ is any robust estimator of scale (e.g. median absolute deviation ($MAD$) or $Qn$, see Han *et al.*, 2019) that quantifies the statistical dispersion of the mean pairwise patristic distance distribution for the ensemble of $N$ subtrees.

Both PhyCLIP and Phydelity include a robust estimator of scale when calculating the respective within-cluster and maximal patristic distance limit to negate assumptions of symmetry in the pairwise distribution of patristic distances. However, PhyCLIP multiplies the robust estimator of scale with an input parameter $\gamma$, which allows the user to calibrate the resolution of clustering to their research question. PhyCLIP also parameterises the inter-cluster divergence limit of clusters in the model using either the Kolmogorov Smirnov test or Kuiper's test. These tests test the null hypothesis that the pairwise sequence distance distributions of every combinatorial pair of putative clusters is empirically equivalent to that if the two subtrees were clustered together.

Statistical significance is inferred if the multiple-testing corrected p-value for the cluster pair is below a user-input false discovery rate.

PhyCLIP therefore has three user-defined parameters: 1. A minimum cluster size. 2. The $\gamma$ multiple of the robust estimator of scale and 3. the false discovery rate for the Kolmogorov Smirnov/Kuiper test. Accordingly, PhyCLIP's parameters need to be calibrated across a range of input values, optimising the global statistical properties of the clustering results to select an optimal parameter set. The optimisation criteria (see Han *et al.*, 2019) are prioritised by the research question, as the clustering resolution and cluster definition are dependent on the question, and therefore the degree of information required to capture ecological, epidemiological and/or evolutionary processes of interest. By comparison, Phydelity only has a single user-defined parameter ($k$) to define $\mathcal{D}_k$ which is optional and can be automatically scaled.

Unlike for Phydelity, PhyCLIP's designated clusters should not be interpreted as sequences linked by rapid transmission events. The use of the global patristic distribution in PhyCLIP sets the within-cluster limit to a resolution too low to only capture homogenous subpopulations of highly similar viruses descendant from the same source in rapid transmission chains. Phydelity's local maximal patristic distance limit was formulated to allow for the identification of these highly similar sequences putatively linked by transmission events.

Phydelity also does not include PhyCLIP's formulation of an inter-cluster divergence test. PhyCLIP's approach is underpowered to detect statistically distinct distributions in phylogenetic trees that do not include a high number of so-called "false-positives" – sequences that are markedly from a different or outgroup cluster. PhyCLIP's approach works well when representative or relatively complete phylogenies are clustered, as there are many outgroup clusters to test against. However, it is unlikely that many non-outbreak sequences will be included in the background diversity of transmission dynamic studies for which Phydelity was formulated. Resultantly, Phydelity relies on distal-dissociation and the local maximal patristic distance limit to set rigorous boundaries on clusters to ensure robust inter-cluster divergence.

### Supplementary Figures

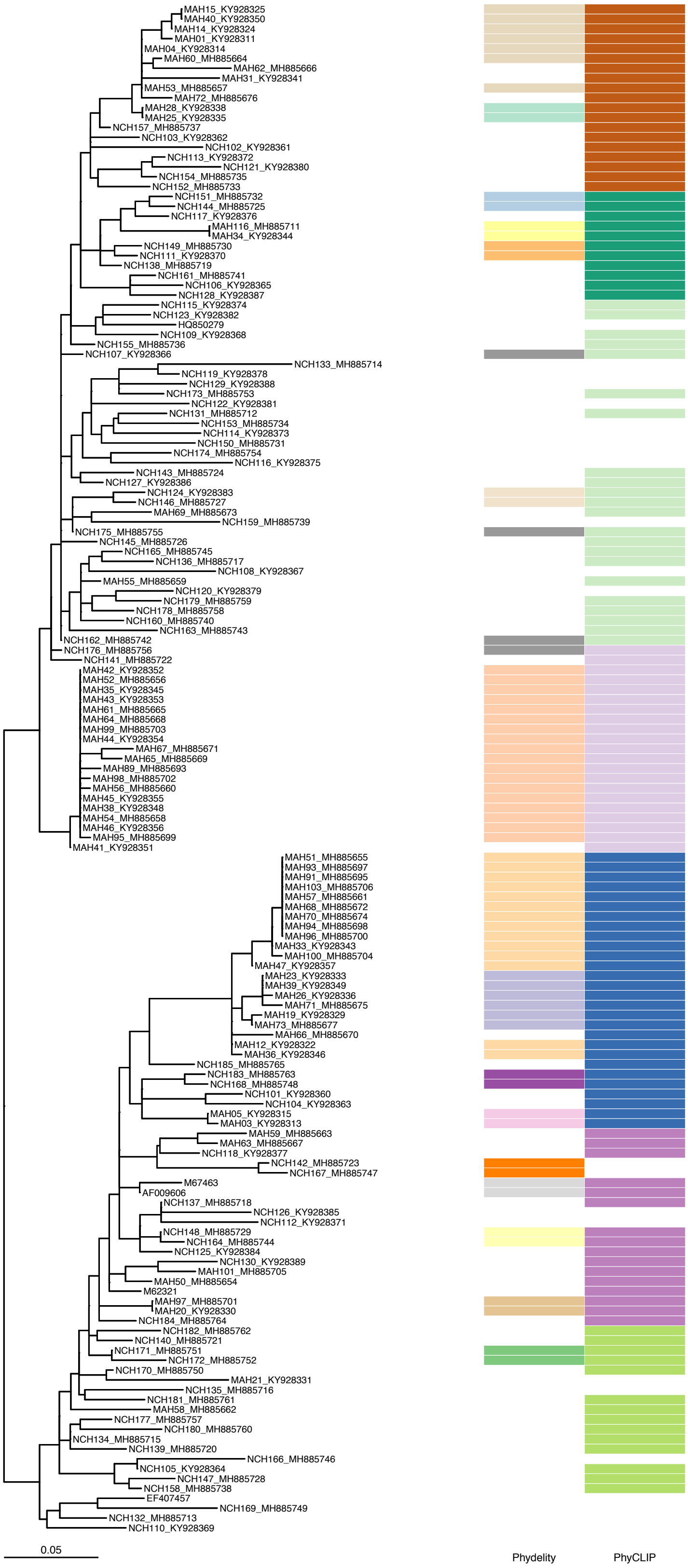


**Supplementary Figure 1**. Comparison of output clusters from Phydelity and PhyCLIP when applied to the Hepatitis C virus genotype 1a NS5B phylogeny. Each distinct colour represents a unique cluster. Input parameters for PhyCLIP are minimum cluster size = 2, $\gamma$ multiple of the robust estimator of scale = 1, and false discovery rate for the Kuiper's test = 0.05.

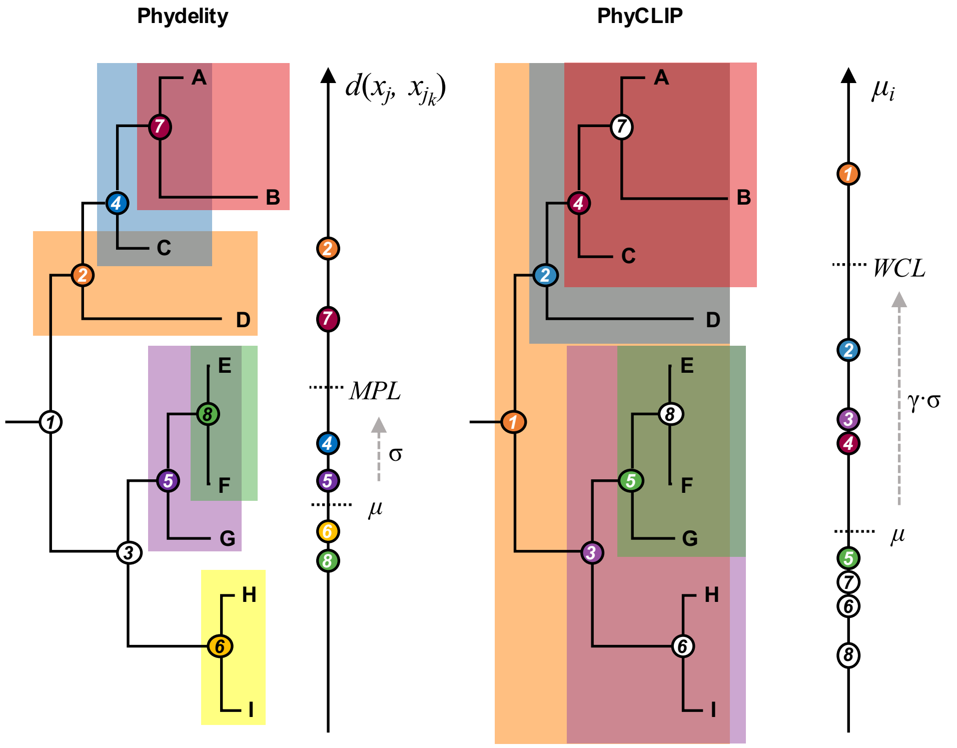


**Supplementary Figure 2**. Comparison between how maximal patristic distance limit (*MPL*) and within-cluster limit (*WCL*) are determined. *MPL* is computed using the median ($\mu$) and robust estimator of scale ($\sigma$) based on the $k$-th core distance distribution ($\mathcal{D}_k$) of every sequence $x_j$ and its $k$-closest neighbours ($d(x_j, x_{j_k})$; $k$=2 in this case). In comparison, PhyCLIP uses the distribution of mean pairwise patristic ($\mu_i$) distance for all putative clusters to calculate *WCL* for a user-provided multiple ($\gamma$) of $\sigma$.

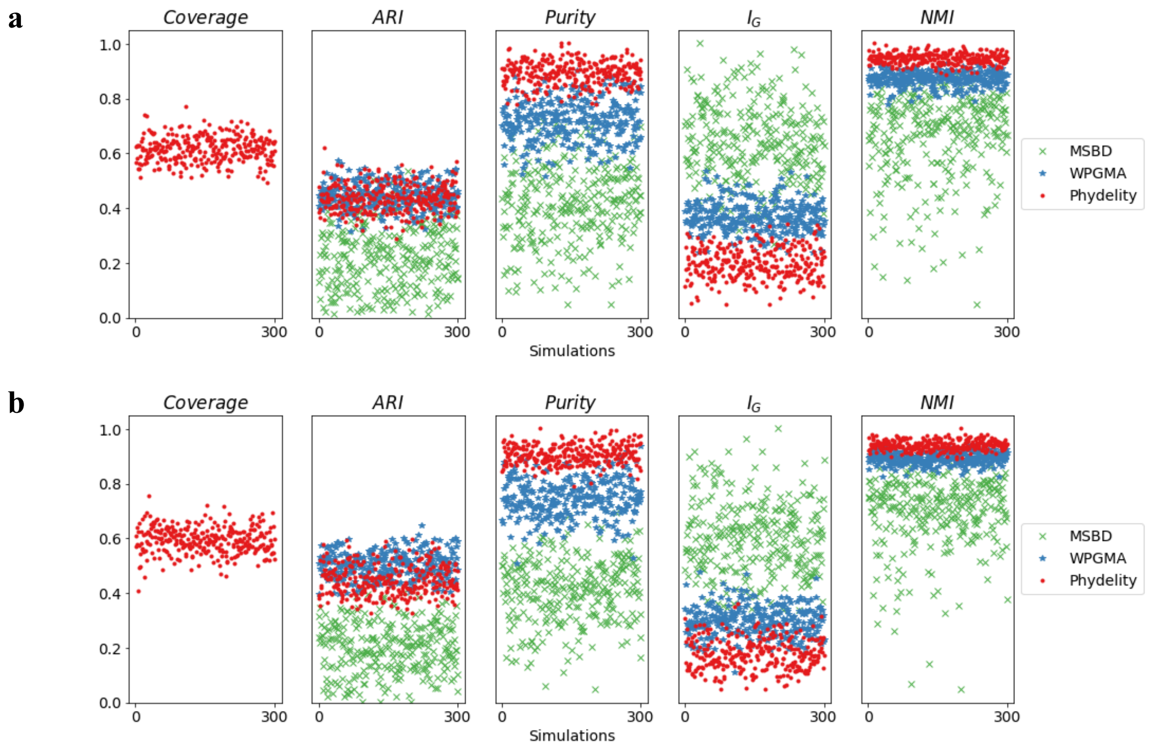


**Supplementary Figure 3**. Clustering results of simulated HIV epidemics in a hypothetical MSM sexual contact network. **(a)** Clustering metrics for clustering algorithms (Phydelity, weighted pair-group method of analysis (WPGMA) and multi-state birth death (MSBD) methods) applied simulated phylogenies with inter-communities transmission rates weighted at half of within-community rates (i.e. $w = 0.25$). Coverage refers to the proportion of all tips clustered (No results shown for MSBD and WPGMA as they clustered 100% of all tips). Adjusted rand index (*ARI*) measures how accurate the output clusters corresponded with the community labels. *Purity* gives the average extent clusters contain only a single class of community. Modified Gini index ($I_G$) is the probbility that a randomly selected sequence would be incorrectly clustered. Normalised mutual information (*NMI*) accounts for the tradeoff between clustering quality and number of clusters. **(b)** Results for simulations where inter-communities transmission rates were identical to within-community rates (i.e. $w = 0.75$).

**a**

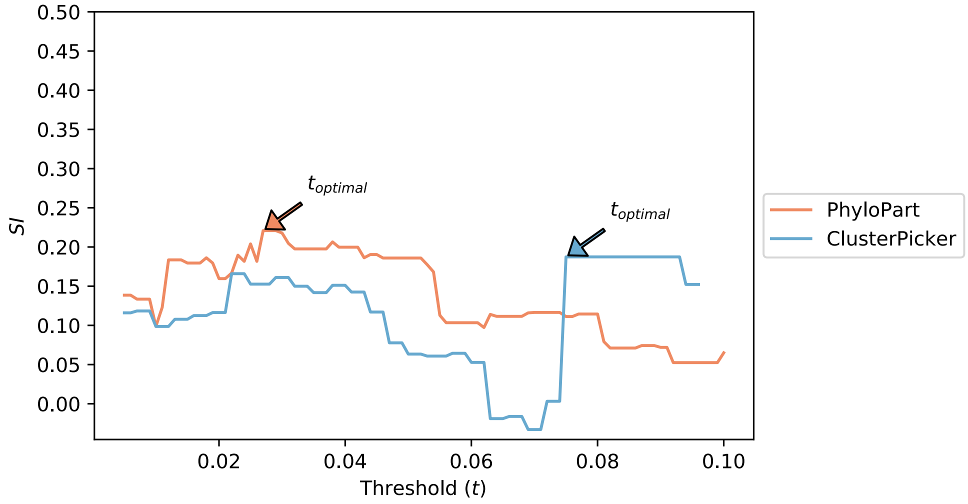


**b**

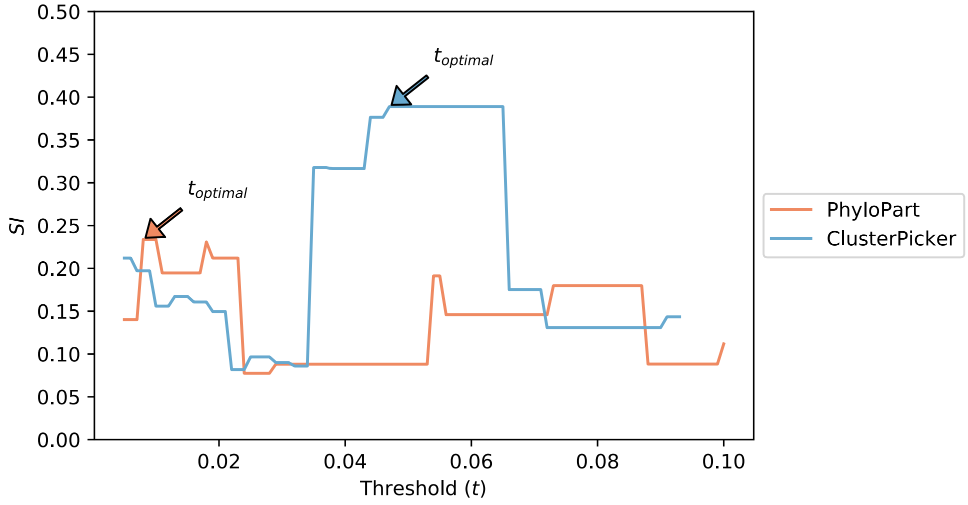


**Supplementary Figure 4**. Plots of mean Silhouette index (*SI*) computed for the range of genetic distance thresholds implemented in ClusterPicker (measured as nucleotides/site) and PhyloPart (measured as percentile of pairwise patristic distribution of input phylogeny). Clustering results from the lowest optimal distance threshold ($t_{optimal}$) with the highest SI value for each method were compared to Phydelity (see Figure 4 in main text). **(a)** HCV genotype 1a ($t_{optimal}$: 0.075 nucleotide/site (ClusterPicker); 2.7% (PhyloPart)). **(b)** HCV genotype 4d ($t_{optimal}$: 0.047 nucleotide/site (ClusterPicker); 0.8% (PhyloPart)).

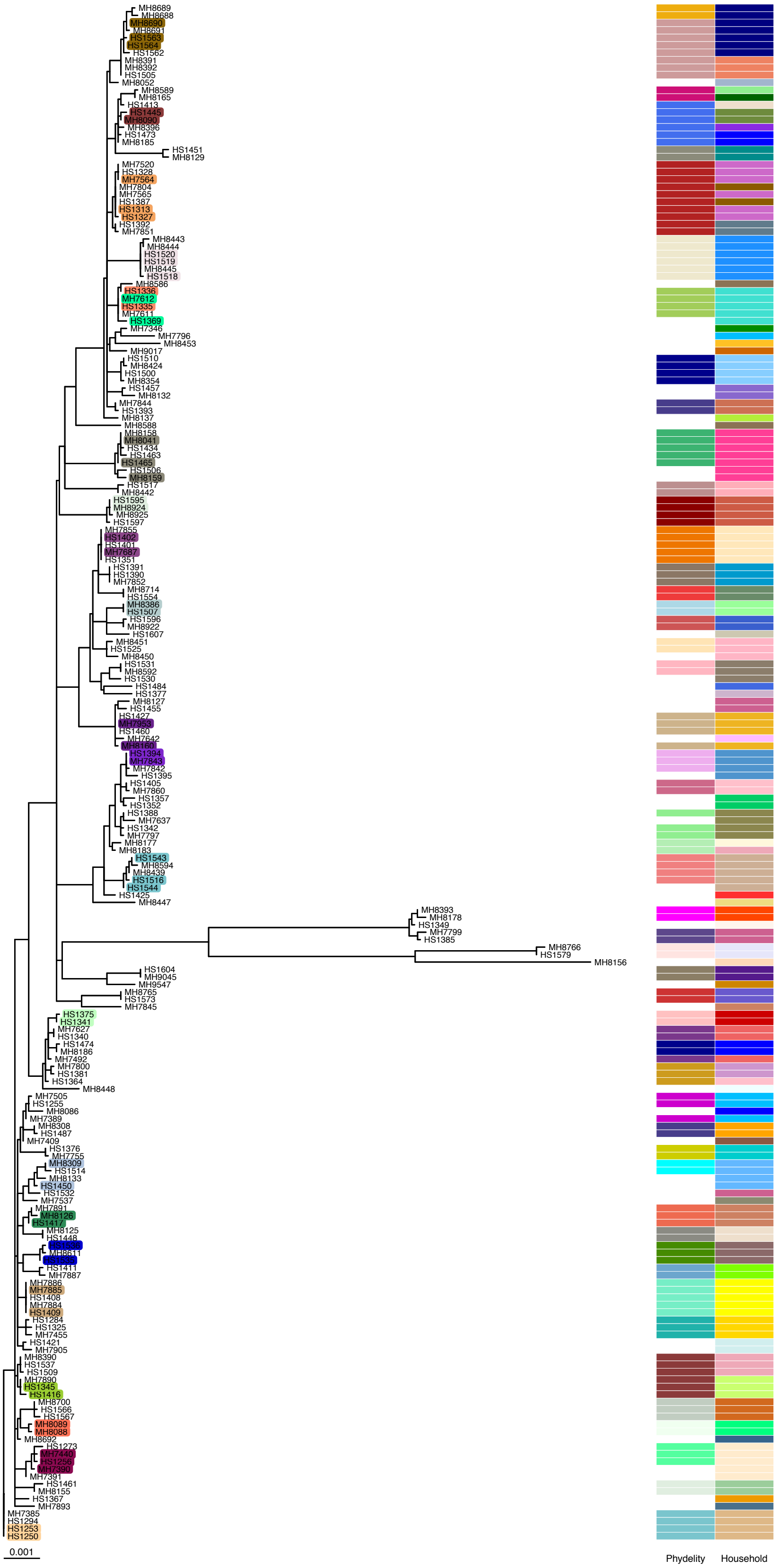


**Supplementary Figure 5**. Phylogenetic tree and Phydelity clustering results of all A/H3N2 influenza viruses collected McCrone et al. (2018). Each distinctly-coloured, highlighted tips represent a different genetically-validated transmission cluster defined by McCrone et al. Clustering results from Phydelity and the household identities of all sequences are depicted as heatmaps. Each distinct colour of the heatmap cells denotes a different Phydelity cluster or household. Note that colours that are similar between Phydelity and household heatmap cells as well as to the highlight tip colours do not imply any form of correspondence.

### Supplementary Tables

| Method | Coverage (SD) | ARI (SD) | Purity (SD) | $I_G$ (SD) | NMI (SD) |
|---|---|---|---|---|---|
| *w = 0.25* | | | | | |
| Phydelity | 61.6% (4.7%) | 0.44 (0.05) | 0.81 (0.03) | 0.27 (0.04) | 0.86 (0.01) |
| WPGMA | N.A. | 0.44 (0.05) | 0.67 (0.06) | 0.40 (0.04) | 0.81 (0.02) |
| MSBD | N.A. | 0.22 (0.11) | 0.43 (0.11) | 0.60 (0.10) | 0.64 (0.11) |
| *w = 0.50* | | | | | |
| Phydelity | 61.2% (4.6%) | 0.44 (0.05) | 0.82 (0.03) | 0.27 (0.04) | 0.86 (0.02) |
| WPGMA | N.A. | 0.47 (0.05) | 0.68 (0.06) | 0.39 (0.04) | 0.81 (0.02) |
| MSBD | N.A. | 0.22 (0.10) | 0.43 (0.10) | 0.59 (0.09) | 0.65 (0.11) |
| *w = 0.75* | | | | | |
| Phydelity | 59.4% (4.9%) | 0.45 (0.05) | 0.85 (0.03) | 0.27 (0.04) | 0.86 (0.02) |
| WPGMA | N.A. | 0.50 (0.05) | 0.71 (0.06) | 0.36 (0.04) | 0.82 (0.02) |
| MSBD | N.A. | 0.20 (0.09) | 0.41 (0.10) | 0.61 (0.10) | 0.64 (0.12) |
| *w = 1.00* | | | | | |
| Phydelity | 58.2% (4.6%) | 0.44 (0.05) | 0.88 (0.03) | 0.28 (0.05) | 0.86 (0.02) |
| WPGMA | N.A. | 0.56 (0.05) | 0.74 (0.06) | 0.33 (0.05) | 0.83 (0.02) |
| MSBD | N.A. | 0.19 (0.09) | 0.39 (0.09) | 0.62 (0.09) | 0.63 (0.10) |

**Supplementary Table 1**. Summary statistics of clustering metrics (mean values and corresponding standard deviation (SD)) comparing results generated from Phydelity, WPGMA and MSBD-method. 300 simulations were performed for each distinct weight ($w$) of inter-community transmission rate. Metrics compared included coverage (i.e. percentage of sequences clustered), adjusted rand index (ARI), purity, modified Gini index ($I_G$) and normalised mutual information (NMI).
